# Increased evolutionary rate in the Z-chromosome of *Morpho* butterflies and implications for speciation

**DOI:** 10.1101/2024.02.02.578590

**Authors:** Manuela López Villavicencio, Joséphine Ledamoisel, Céline Lopez-Roques, Vincent Debat, Violaine Llaurens

## Abstract

The evolution of reproductive isolation between populations shapes divergence in genome structure and content: comparing the genomes of closely-related species can thus enlighten the speciation process. Comparisons of genomes of allopatric *vs*. sympatric species sharing similar *vs*. dissimilar ecological niches allows to specifically investigate the effect of reinforcement and ecological specialization on genome evolution. In the butterfly genus *Morpho*, several species can be found in sympatry presenting specialisation in different microhabitats and temporal niches. Here, we sequenced, assembled and annotated the genomes of 8 *Morpho* species and used previously published genomes of three other *Morpho* species to study genomic rearrangements and signatures of positive selection. We found extensive chromosomal rearrangements in the Z chromosome compared to the autosomes, particularly among closely related sympatric species occupying similar niches, pointing at the putative role of inversions in preventing gene flow at a postzygotic level. We also detected a higher proportion of genes under positive selection on the Z-chromosome compared to the autosomes, suggesting a potential role of the Z-chromosome in driving adaptive evolution in *Morpho*. Finally, because of the divergence in daily activities between species, we studied the evolution of eight genes involved in the circadian clock and detected a signature of positive selection on the gene *Period*, located in the Z chromosome. By studying the evolution of genome structure and coding sequences, our study indicates fast evolution of the Z-chromosome, partly driven by selection, throughout this genus, highlighting the putative implication of this sexual chromosome on pre and post-zygotic isolation.

## Introduction

Ecological and genomic mechanisms involved in the process of speciation are still largely unknown. Identifying factors increasing pre and post zygotic isolation between populations is especially important to understand how species can diverge and co-exist in sympatry. In sympatry, specialisation into different ecological niches may strongly limit gene flow between species because of (1) reduced encountering rates between individuals specialised on different niches and (2) decreased fitness of hybrids that may be less adapted to either niches [55]. The genomic factors underlying such pre- and post-mating isolation are extensively studied in multiple taxa, and divergence in the sex chromosome (X or Z depending on the considered species) has been shown to play a significant role [57]. The divergence in Z-chromosome is assumed to result in poor hybrid fitness, acting as a post-zygotic barrier to gene flow between diverging populations. In passerine birds for instance, chromosomal inversions were observed to be more prevalent on the Z-chromosomes as compared to the autosomes, and interestingly the different gene order was more likely to be fixed within species, in species pairs with overlapping geographic distribution, suggesting an effect of reinforcement [33].

Furthermore, genes involved in sexual dimorphism and mate preferences are also frequently found on the sex chromosomes and may also contribute to pre-mating isolation: for instance, in closely-related species of *Fiducela* flycatchers, loci controlling plumage divergence and preference for plumage coloration are located on the Z-chromosomes, and are likely to enhance pre-mating barriers [63]. In Lepidoptera, sex determination also stems from ZZ/ZW sex chromosomes, and evidence for a significant role of the Z-chromosome in speciation has been reported. In a pair of sister-species of *Colias* butterflies with overlapping geographic range, increased genetic divergence in the Z-chromosomes as compared to the autosomes was observed. Moreover, loci controlling for divergent coloration in males are located on the Z-chromosome, suggesting a key role of the Z-chromosome in the pre-zygotic barrier to gene flow in these species [23].

Here, we focus on the genomic divergence between species of *Morpho* butterflies, where multiple species currently co-exist in sympatry in the Amazonian rainforest [4]. In the genus *Morpho*, sympatric species show divergence in (1) flight height, with specialisation in the canopy *vs.* understory microhabitats [40] and in (2) flight time [41], with males from some species only flying early in the morning while males from other species typically patrol at noon. Such specialisation into different forest strata and/or temporal niches likely contributes to the co-existence of these closely-related species in sympatry and may also have played a role in the speciation process itself. In the noctuid moth *Spodoptera frugiperda*, populations specialised on different host-plants diverge in the timing of mating, and the gene *vrille*, involved in the control of the circadian clock, was shown to diverge between these species [29]. The evolution of circadian genes may thus be involved in the divergence of temporal niches, therefore enhancing pre-zygotic barrier to gene flow in sympatric species. Interestingly, in Lepidoptera, some genes involved in the regulation of circadian behaviour are sex-linked: for instance the gene *Period*, responsible for diurnal or nocturnal activity in some butterflies and moths or the gene *Clock*, involved in the photoperiodic response and migration in the Monarch buterfly, are both located on the Z-chromosome [28, 34, 70].

We report the sequencing, assembly and annotation of the genomes of 8 species from the genus *Morpho*. We then used these new genome assemblies, as well as the previously published genomes of 3 other *Morpho* species [3] to perform comparative genomic analyses in 11 out of the 30 species of the genus *Morpho*. We specifically compare the evolution of the Z-chromosome *vs*. the autosomes, in sympatric and allopatric *Morpho* species, specialized in different ecological niches: we investigate the density of repeated elements, inversions and genes under positive selection in the Z-chromosome *vs.* the autosomes. We also investigate the location and evolution of eight genes known to regulate circadian activities in other Lepidoptera (*Clock, Cycle, Period, Timeless, Cry1, Cry2, Vrille and PDP1*) [6, 60], and specifically test for signature of positive selection on these genes.

## Materials and Methods

The nuclear and mitochondrial assemblies of the genomes of *Morpho helenor*, *M* . *achilles* and *M* . *deidamia* were previously described in [3]. Here, we sequenced, assembled and annotated the whole genome of eight additional *Morpho* species: five understory species - *M* . *amathonte*, *M* . *menelaus*, *M* . *granadensis*, *M* . *eugenia* and *M* . *marcus* - and three canopy species - *M* . *rhetenor*, *M* . *hecuba* and *M* . *telemachus*. For this last species, two different morphs co-exist in sympatry: males with grey or golden wings, and we sequenced one individual of each morph.

### Butterfly sampling

The male butterflies used for sequencing in this study were collected with handnets in Panama, Veraguas district (GPS coordinates: 8.525869707300163, -81.13020529212375) in July 2022 for *M. granadensis* and *M. amathonte* (Export permit number:

PA-01-ARB-076-2022) and in French Guiana, Roura district (GPS coordinates: 4.5917807378071975, -52.20357910219149) in october 2022 for *M. eugenia, M. marcus, M. hecuba, M. telemachus* grey and gold morphs, *M. menelaus, M. rhetenor* (authorization number: R03-22-10-18-0001). The butterfly samples were then SNAP-frozen and kept at -80°C.

### DNA extractions and genome sequencing

For genomic analyses, the DNA extraction was carried out from the thorax muscles using the Qiagen Genomic-tip 100/G kit, following supplier instructions. Library preparation and sequencing were performed according to the manufacturer’s instructions “Procedure & Checklist – Preparing whole genome and metagenome libraries using SMRTbell® prep kit 3.0 ». At each step, DNA was quantified using the Qubit dsDNA HS Assay Kit (Life Technologies). DNA purity was tested using the nanodrop (Thermofisher) and size distribution and degradation assessed using the Femto pulse Genomic DNA 165 kb Kit (Agilent). Purification steps were performed using AMPure PB beads (PacBio) and SMRTbell cleanup beads (PacBio). For 9 samples, 5µg of DNA was purified then repaired using “SMRTbell Damage Repair Kit SPV3” (PacBio) and sheared between 15kb to 20 kb using the Megaruptor 1 system (Diagenode). For *Morpho amathonte*, *M* . *granadensis* and *M* . *telemachus* gold samples, a single library was prepared. For the other 6 samples, multiplex libraries were prepared using SMRTbell Barcoded Adaptater kit 8A for *M* . *hecuba*/*M* . *rhetenor*, *M* . *telemachus* grey/*M* . *marcus* and *M* . *eugenia*/*M* . *menelaus*. A nuclease treatment was performed on the 6 libraries folowed by a size selection step using a 6, 8 or 10kb cut off on the PippinHT Size Selection system (Sage Science) with “0.75% Agarose, 6-10 kb High Pass, 75E protocol“. Using Binding kit 3.2 kit and sequencing kit 2.0, libraries were sequenced with the adaptive loading method onto 6 SMRTcells (+2 extra SMRTcells for *M* . *menelaus* sample) on Sequel2 instrument at 90 pM with a 2 hours pre-extension and a 30 hours movie.

### Nuclear and mitochondrial genome assembly

Heterozygosity for each PacBio dataset was estimated with Jellyfish (v.2.3.0) [47] (jellyfish count and hist options). The resulting histograms were visualized with GenomeScope online version 2.0 [59]. Assemblies were performed using Hifiasm v0.19.5 [15]. Lepidoptera genomes are particularly heterozygous and high heterozygosity imposes a challenge for most genome assemblers as it may be difficult to make the distinction between different alleles at the same locus and paralogs at different loci.

Raw assemblies from heterozygous genomes are thus expected to contain high levels of false duplications [2]. Hifiasm allows using different levels of purge to remove redundant contigs. We systematically produced assemblies using two different purge options: no purge and purge in the most aggressive way (options -l0 and -l3 respectively). For each assembly, we estimated basic statistics (number of contigs, size of contigs, N50) using the “stats.sh” program from the BBMap v38.93 package [9]. We assessed the completeness of assemblies obtained using the different options with BUSCO v5.2.2 and MetaEuk for gene prediction against the *lepidoptera*_*odb*10 database [46]. In several cases, the BUSCO results showed a high level of duplications in the final assembly, even when using the aggressive mode in hifiasm. In these cases, we used Purge_dups v1.2.5 to remove false haplotypic duplications, and then we reassessed completeness using BUSCO. For every species, we chose the final assembly among the different assemblies produced with or without the use of Purge_dups, based in the basic statistics and the BUSCO score. The mitochondrial genome for all *Morpho* species was assembled directly from the PacBio Hifi reads with Rebaler (https://github.com/rrwick/Rebaler) as described in [3] and using the mitochondrial genome from *M* . *helenor* as a reference.

All sequences obtained in this study have been submitted to NCBI GenBank.

### Genome annotation and identification of circadian genes

For each *Morpho* species, we created a *de novo* library of repetitive elements from each assembled genome with RepeatModeler v2.0.2a [24] with the option -s (slow search). This library was then used with RepeatMasker 4.1.2.p1 [24] to produce softmasked assemblies needed for the annotation.

Structural annotation was carried out with BRAKER v2.1.6 [7, 31], using the *arthropoda*_*odb*10 protein data set from OrthoDB [39]. BRAKER2 *ab initio* gene predictions were performed with AUGUSTUS v3.4.0b [8] and GeneMark-EP+ v4.64 [43] with two rounds of training. The resulting .gff files containing predicted genes were filtered to keep only the longest isoform and were screened to remove incomplete gene coding models (i.e. genes coding with start and/or stop codon missing in its cds) with AGAT [17]. The filtered .gff files produced were used in the following steps.

The functional annotation was added to the BRAKER2 predictions in two parts, first, we ran InterProScan v5.33-72.0 [58] on the files produced after two rounds of BRAKER2 using the option -f XML. Second, the XML file produced by InterProScan and the previously filtered .gff file were used in Funannotate v1.8.13 [56] with the option “annotate” to obtain a multi-fasta file of protein coding genes for each species. We followed this procedure on the nine new genomes assembled in this paper and on the three previously published genomes of *M* . *helenor*, *M* . *achilles* and *M* . *deidamia*, to obtain comparable annotations throughout the 12 genomes. To assess the quality of the gene-set predicted for each species, we ran BUSCO analyses on the protein file produced selecting the protein mode and the *lepidoptera*_*odb*10 database.

We then specifically studied eight different genes known to be involved in the circadian clock in butterflies: *Clock, Cycle, Period, Timeless, Cry1, Cry2, Vrille* and *PDP1*). The position, the proteic and nucleotidic sequences of the genes were extracted from the final gff files produced at the annotation step.

### Phylogenetic relationships between ***Morpho*** species

Phylogenetic analyses were conducted separately for nuclear and mitochondrial DNA data sets. To infer the phylogenetic relationships between the 12 sequenced genomes of *Morpho* butterflies, we followed the procedure explained in [72]. Single copy orthologs for each genome were identified from BUSCO proteome results using busco2fasta.py (https://github.com/lstevens17/busco2fasta) with the protein option. Orthologs were aligned using Mafft with default options [38] and trimmed using trimAl [12]. Trimmed alignments were concatenated to produce a phylip supermatrix with catfasta2phyml (https://github.com/nylander/catfasta2phyml). This matrix was used in IQ-TREE [53] to infer the species tree under the JTT+F+R10 model chosen by ModelFinder using tree search [37], SH-aLRT test and ultrafast bootstrap with 1000 replicates [30]. We follow the same procedure with IQ-TREE to construct the mitochondrial tree of the group using the mitochondrial sequences previously obtained with Rebaler. The consensus trees were visualized using FigTree v1.4.4 (http://tree.bio.ed.ac.uk/software/figtree/).

### Synteny and detection of chromosomal rearrangements between ***Morpho* genomes**

To study the synteny and major rearrangements, we aligned the genomes of closely-related species by pair, using the online version of D-genies [10]. Given our focus on synteny among closely related species, we conducted comparisons involving all species within the “Canopy” clade. Specifically, three comparisons were performed: *Morpho rhetenor* vs *M* . *hecuba*, *M* . *hecuba* vs *M* . *telemachus* grey, and both morphs of *M* . *telemachus*. For the “Understory” clade, we also performed comparisons between close species, specifically: *M* . *amathonte* vs *M* . *menelaus*, *M* . *granadensis* vs *M* . *helenor* and *M* . *helenor* vs. *M* . *achilles*. Finally, we added the paired analysis of the two basal sister species: *M* . *eugenia* vs *M* . *marcus*. Paired alignments were performed using the Minimap2 aligner [42] in D-genies (see Fig. S4). The contig corresponding to the Z chromosome for *M. helenor*, *M* . *achilles* and *M* . *deidamia*, were already identified [3]. We then used D-genies to pair-map the genome of *M. helenor* to the newly assembled *Morpho* species genome (*M* . *rhetenor*, *M* . *hecuba*, *M* . *telemachus*, *M* . *amathonte*, *M* . *menelaus*, *M* . *granadensis*, *M* . *eugenia* and *M* . *marcus*) treating *M. helenor* genome as the target reference and the new *Morpho* species genome as the query. This alignment permitted us to visually find, for each species, the contig corresponding to the already known Z chromosome of *M. helenor*.

In our paired whole-genome comparisons, inversions within the contig corresponding to the Z chromosome were recurrently observed across various species (see Fig. S4).

Subsequently, we employed SyRI for targeted paired comparisons, specifically concentrating on the Z chromosome between the same closely-related species pairs used for whole-genome comparisons. Paired comparisons followed the methodology described in [3]. Briefly, paired alignments of the Z contig were generated using Minimap2, and the resulting SAM files were analyzed with SyRI to detect inversions [27]. The genomic structures predicted by SyRI were visually represented using plotsr [26].

Repeated elements are expected to accumulate in sexual chromosomes due to reduced recombination rate as compared to autosomes [22]. Repeated elements are also assumed to promote chromosomal inversions, since they are frequently found prevalent at the margin of inversion breakpoints (*e.g.* in Drosophila, [11]). We thus assessed whether the contigs corresponding to the Z chromosome in different *Morpho* species exhibited a higher frequency of repeated elements as compared to the rest of the contigs in the genome. We identified repeated regions using RepeatModeler v2.0.2a [24] and RepeatMasker [24] separately for each contig within the different genome assemblies. Because SyRI plots showed that inversions between paired *Morpho* species were usually found at the margins of the contig Z, we looked for potential alterations in the distribution of repeated elements along this contig. For each species, we partitioned the Z contig into approximately 5Mb sections and conducted RepeatModeler and RepeatMasker analyses for each section. This procedure was replicated in two additional autosomal contigs of approximately equivalent sizes for comparison. We discarded contigs spanning less than 10Mb from the assemblies, to simplify subsequent analyses.

### Detection of positive selection in circadian genes and throughout the whole genome

To explore the genetic basis of the divergence in temporal niches, we searched for signature of selection in eight circadian genes. We first generated trees for each gene with IQ-TREE [53] and ModelFinder. These trees were then used in the Codeml program in the PAML package (version 4.8) [1, 74] to detect signals of positive selection. PAML estimates the omega ratio (*dN/dS*) using maximum likelihood. The omega ratio compares non-synonymous (*dN*) against synonymous (*dS*) substitutions per site. We used random site models to test for positive selection at the codon scale on the genes. Branch site models were also computed to estimate selective pressures at codon sites within different lineages. Three types of hypotheses were tested under the PAML branch site model: (1) we searched for differences of positive selection on the branch leading to the two most basal *Morpho* species: *M.marcus* and *M.eugenia vs.* as this clade was the earliest diverging clade in the genus around 28MYA [13]. We labeled these two species as foreground branches and the rest of the species as background branches; (2) as the evolution of circadian genes might be linked to differences in the amount of light available, we looked for traces of positive selection on the lineages of the *Morpho* species living in the canopy *vs* the understory; and (3) because time activity is expected to be influenced by circadian genes, we looked for evidence of positive selection in the species where males display an early flight activity (patrolling from 6:00 to 10:00 am) *vs* a late flight activity (patrolling after 10:00 am). All models were always compared to a null model assuming an omega ratio of 1 for all sites and across all lineages. To determine significance, the null model and the positive selection model were compared using a likelihood ratio test (LRT) following a Chi2 distribution, and Bayes Empirical Bayes (BEB) analyses were run to identify the positively selected amino acids [1, 74]. All the analyses were carried out using PAMLX [73]

To detect differential selection in protein sequence alignments at the genome-wide level, we then used the software *Pelican* [21]. Based on a maximum likelihood approach, *Pelican* aims at detecting significant shifts in amino acid preferences between lineages. The software scans alignments for sites differing in amino acid preferences depending on a phenotypic trait that is specified on a phylogenetic tree. We used the single copy ortholog alignments found in all the 12 genomes sequenced and described in the section “*Phylogenetic relationships between Morpho species*“. To estimate differences in selection regime in the different micro-habitats (canopy or understory), we used two different trees: a first tree (“Canopy”) labeling the canopy species in foreground (trait=1) and the understory species plus the basal clade labeled as background (trait=0) and a second tree (“Understory”) with the clade containing the understory species labeled as foreground and the canopy and basal species as background. We used the option pelican scan discrete and –alphabet=AA. *Pelican* produces a file containing one *p-value* per site across all genes and assigns a *p-value* equal 1 to amino acids that are shared across all species [20]. Pelican does not produce gene-level predictions directly, nevertheless, the Gene-wise Truncated Fisher’s method (GTF) can be applied to obtain a score for each gene. This method has been shown to be reliable for empirical and simulation data https://gitlab.in2p3.fr/phoogle/pelican/-/wikis/Gene-level-predictions [20].

For each tree, we selected the 50 first candidate genes (genes with the lowest *p-values* produced by *Pelican*), noting the contigs on which they were located. Given that an over-representation of candidate genes for specific contigs may be attributed solely to differences in the gene count within those contigs, we retrieved the genes used for the alignments in *Pelican* to calculate the overall gene count per contig. This assessment was further adjusted based on the contig size, measured as the number of genes per megabase (Mb). We also calculated the gene density by determining the number of genes per megabase in the remaining contigs that exceeded 10 megabases in the genome. This analysis was performed for comparative purposes.

## Results

### Nuclear and mitochondrial genome assembly

Nuclear genome sizes of the Morpho species ranged from 375 to 514Mb (see Table 1). Consistent with other Lepidoptera and with the known genomes of other *Morpho* species, Genomescope analyses showed high heterozygosity for all the species, ranging from 1% of heterozygosity for *M* . *granadensis* to 3.38% for *M. menelaus*. In all the cases, BUSCO results showed a very high level of duplicated sequences, when the assembly was performed using the option -l0 (no purge) in Hifiasm and in most cases, we had to use the option -l3 (most aggressive purge). The BUSCO values then showed low levels of duplicates in the assemblies for *M* . *telemachus*, *M* . *amathonte* and *M* . *granadensis*. In contrast, *M* . *hecuba*, *M* . *menelaus*, *M* . *eugenia* and *M* . *marcus* genome assemblies still showed high levels of duplicates despite the use of the l3 option and needed to be further purged using purge_dups. In the case of *M* . *rhetenor*, the best assembly was obtained without purging in Hifiasm, but with the use of purge_dups. Final BUSCO results are shown in the figure S1 (upper). Mitochondrial genomes presented similar sizes to the previously assembled genomes of other *Morpho* species [3]. For the nine individuals sequenced here, the Rebaler software detected mitochondrial genomes, with sizes ranging from 13,469 bp for the grey morph of *M. telemachus* to 16,173 bp in *M. amathonte*.

**Table 1.**
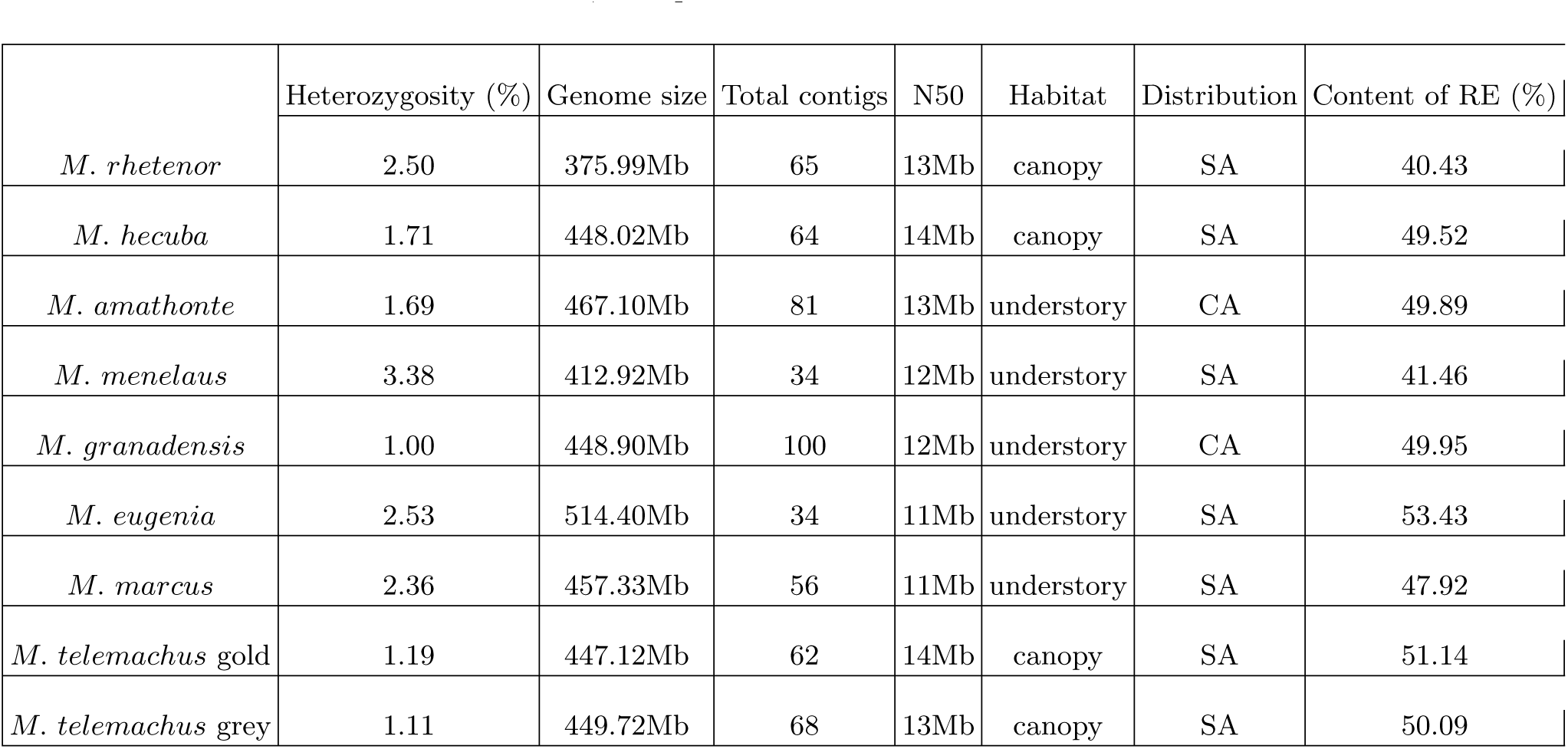
Basic statistic for the assemblies of eight *Morpho* species. Assemblies were performed using Hifiasm and were purged using purge_dups when needed based on BUSCO results on the assembly. The habitat (understory or canopy) and the distribution (Central America - CA or South America - SA) is indicated for each species

|  | Heterozygosity (%) | Genome size | Total contigs | N50 | Habitat | Distribution | Content of RE (%) |
| --- | --- | --- | --- | --- | --- | --- | --- |
| *M. rhetenor* | 2.50 | 375.99Mb | 65 | 13Mb | canopy | SA | 40.43 |
| *M. hecuba* | 1.71 | 448.02Mb | 64 | 14Mb | canopy | SA | 49.52 |
| *M. amathonte* | 1.69 | 467.10Mb | 81 | 13Mb | understory | CA | 49.89 |
| *M. menelaus* | 3.38 | 412.92Mb | 34 | 12Mb | understory | SA | 41.46 |
| *M. granadensis* | 1.00 | 448.90Mb | 100 | 12Mb | understory | CA | 49.95 |
| *M. eugenia* | 2.53 | 514.40Mb | 34 | 11Mb | understory | SA | 53.43 |
| *M. marcus* | 2.36 | 457.33Mb | 56 | 11Mb | understory | SA | 47.92 |
| *M. telemachus* gold | 1.19 | 447.12Mb | 62 | 14Mb | canopy | SA | 51.14 |
| *M. telemachus* grey | 1.11 | 449.72Mb | 68 | 13Mb | canopy | SA | 50.09 |

### Genome annotation

Similarly to results obtained in other *Morpho* genomes, we detected that 40% to 53% of the genomes of the 9 *Morpho* sequenced here consisted of repeated elements (Table 1) By re-annotating the three previously published *Morpho* genomes (*M* . *helenor*, *M* . *achilles* and *M* . *deidamia*), we found significantly more genes predicted and better proteome BUSCO scores using BRAKER2 than with the previous methodology using Maker v2.31.10 [3]. Overall, for most *Morpho* species, we annotated between 18,159 and 21,017 protein coding genes in their genomes (see table S2). These values are close to what has been found using similar procedures in other Nymphalidae species, as *Euphydryas editha* [67] or *Lacanobia w − latinum* [5]. In *M.helenor* however, we found an excess of annotated genes, as we recovered 29,734 complete genes.This elevated number of annotated genes is unlikely to stem from artefactual gene duplications since the BUSCO score in this species did not revealed any increase in gene duplication as compared to the other species.

### Phylogenetic relationships between ***Morpho*** species

We inferred the phylogenetic relationships among samples using 4,333 single-copy orthologs present in all the 12 available genomes of *Morpho*. This phylogeny is consistent with the published phylogeny of the group constructed using mitochondrial and nuclear genes [13, 14]. All the canopy species are grouped together in a single monophyletic clade. Understory species are composed of two clades, a basal clade containing only *M* . *eugenia* and *M* . *marcus* and a bigger clade containing the rest of the understory species. The divergence between the grey and the gold morphs within the species *M. telemachus* was very low, confirming that these two forms found in sympatry indeed belong to the same species.

### Synteny and detection of chromosomal rearrangements between ***Morpho* genomes**

We identified the contigs corresponding to the Z chromosome in all the *Morpho* species. The size of this contig exhibited variation across species, ranging from approximately 18 to 25 Mb. Notably, the Z contig constituted the largest contig in the assemblies of all species, except *M* . *marcus* and *M* . *eugenia*,(see S1). D-genies paired alignments of the whole genome showed strong synteny in all autosomes across the majority of closely-related species (see S4).

When specifically comparing the Z contig among pairs of closely-related species (see S4), three among the seven paired comparisons unveiled variable-sized inversions (Fig 2). Within the canopy clade, a 2.5 Mb inversion was identified between *M. telemachus* and *M* . *hecuba* (Fig 2, subfigure a). While D-genies whole-genome comparisons uncovered two small inversions between *M* . *rhetenor* and *M* . *hecuba* (see S4) these were not observed in the paired comparisons conducted with SyRI (see Fig 2 subfigure a). SyRI plots between these two species revealed lower synteny in the Z contig compared to other species pairs, suggesting that SyRI performs more effectively in genomes with high synteny. For the species living in the understory, SyRI plots revealed three inversions positioned at the tips of the Z contig among the three species *M* . *granadensis*, *M* . *achilles*, *M* . *helenor*(Fig 2 subfigure c). Comparisons also showed two pairs of closely-related species without any inversions and with a very high synteny level along the entire Z-chromosome contig (*M* . *menelaus* vs *M* . *amathonte* and *M* . *eugenia* vs *M* . *marcus* Fig 2 subfigures b and d).

**Figure 1.**
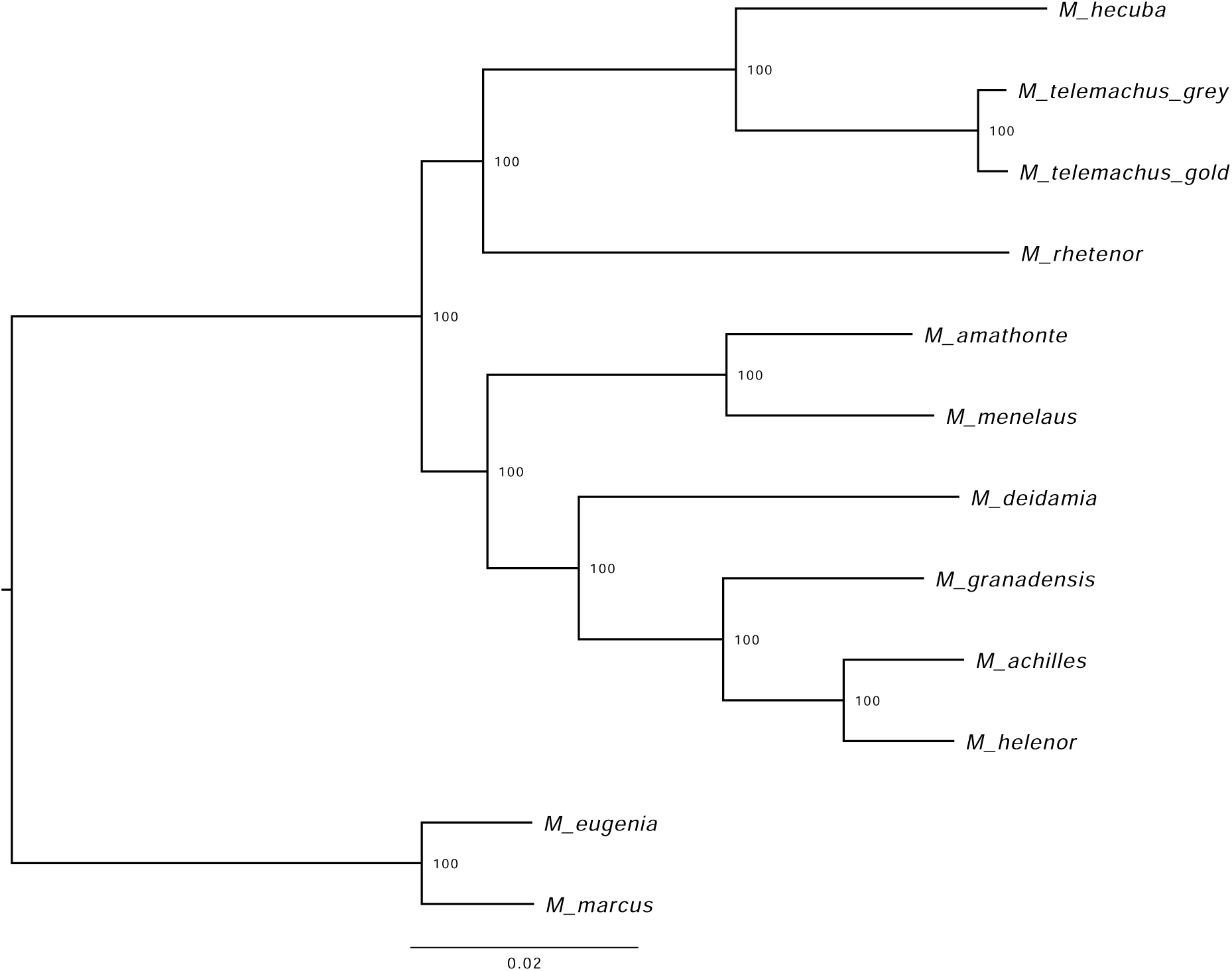
Phylogenetic relationships between the 12 *Morpho* samples. Phylogenetic relationships were inferred using IQ-TREE together with GTR+F+R10 model with a supermatrix from 4,333 single-copy orthologs found in all the samples

**Figure 2.**
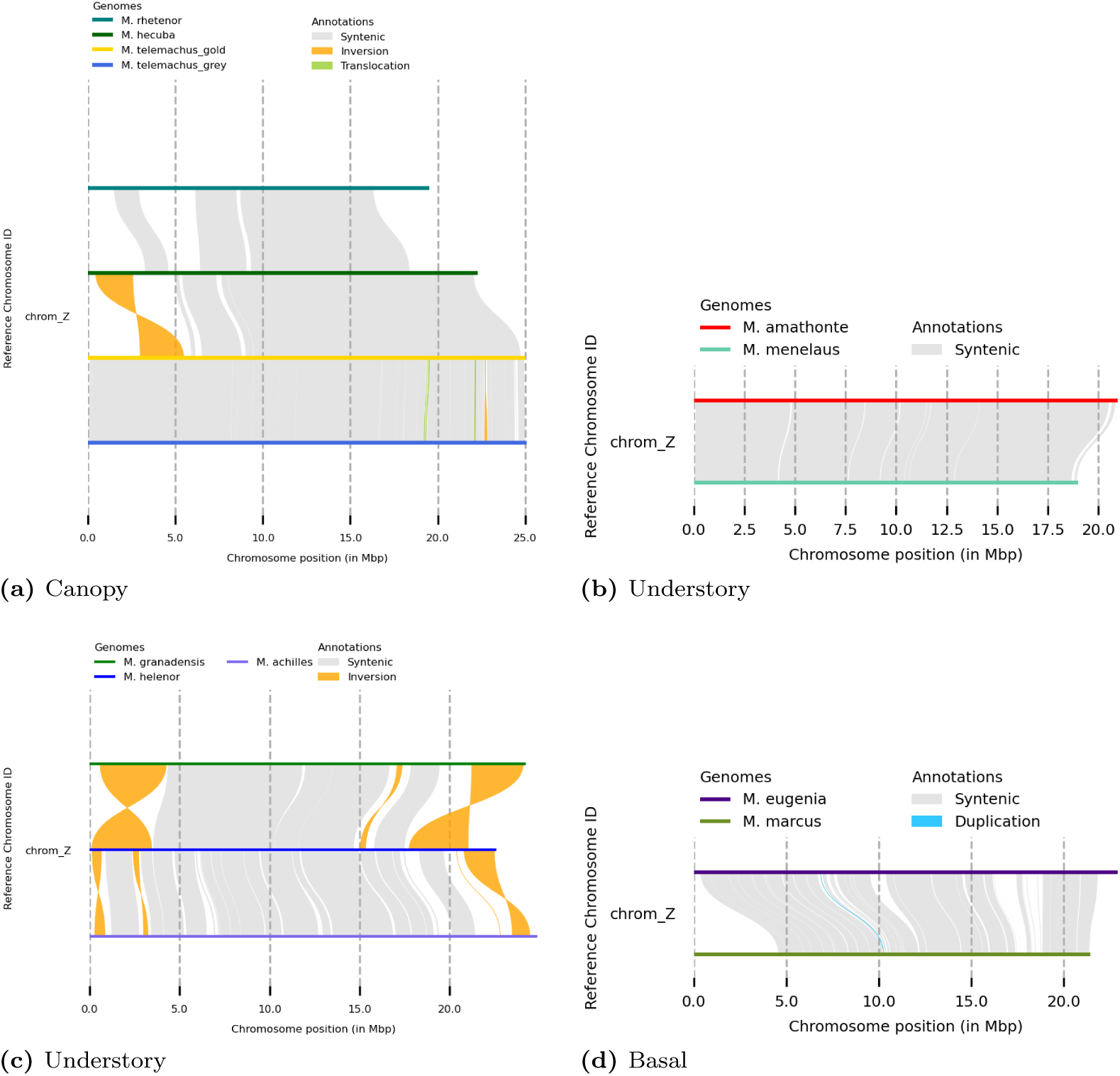
Synteny between the Z chromosome of different *Morpho* species. Pairwise comparisons between closely-related species from a) the canopy clade b) understory clade *M. amathonte* and *M. menelaus* c) understory clade *M. granadensis* and sister species *M. achilles* and *M. helenor* d) basal clade *M. eugenia* and *M. marcus*. Inversions are visible in dark yellow and syntenic regions in light gray repeated elements in the most external sections of the contig, coinciding with regions where inversions typically occur. However, similar results were observed for partitions of contigs corresponding to autosomes where no inversions were observed (see Fig S3).

For all the species, the frequency of repeated elements was not significantly higher in the Z contig as compared to other contigs (see table S2). The analyses of the partitions of the Z contig using RepeatModeler and RepeatMasker revealed a higher content of

This suggests that the accumulation of repeated elements at the chromosomal tips is a shared characteristic across the entire genome, and does not necessarily correlate with increased probability of chromosomal rearrangements.

### Signatures of positive selection in circadian genes

The random site models implemented in PAML applied to our 12 Morpho samples did not reveal any signal of positive selection on any of the eight circadian genes tested (Fig S5). However, using a branch site model, significant evidence of positive selection was found in one of these genes when comparing the canopy vs. understory species: a significant signature of positive selection in the canopy lineage was found in *Period*, located on the Z contig (*p-value*=0.03*, see Fig S6 for analyses details). Such positive selection detected on sites where amino acids are different between the canopy and the understory species suggests that different selective pressures have affected the evolution of the *Period* gene in the two micro-habitats (Fig 3).

**Figure 3.**
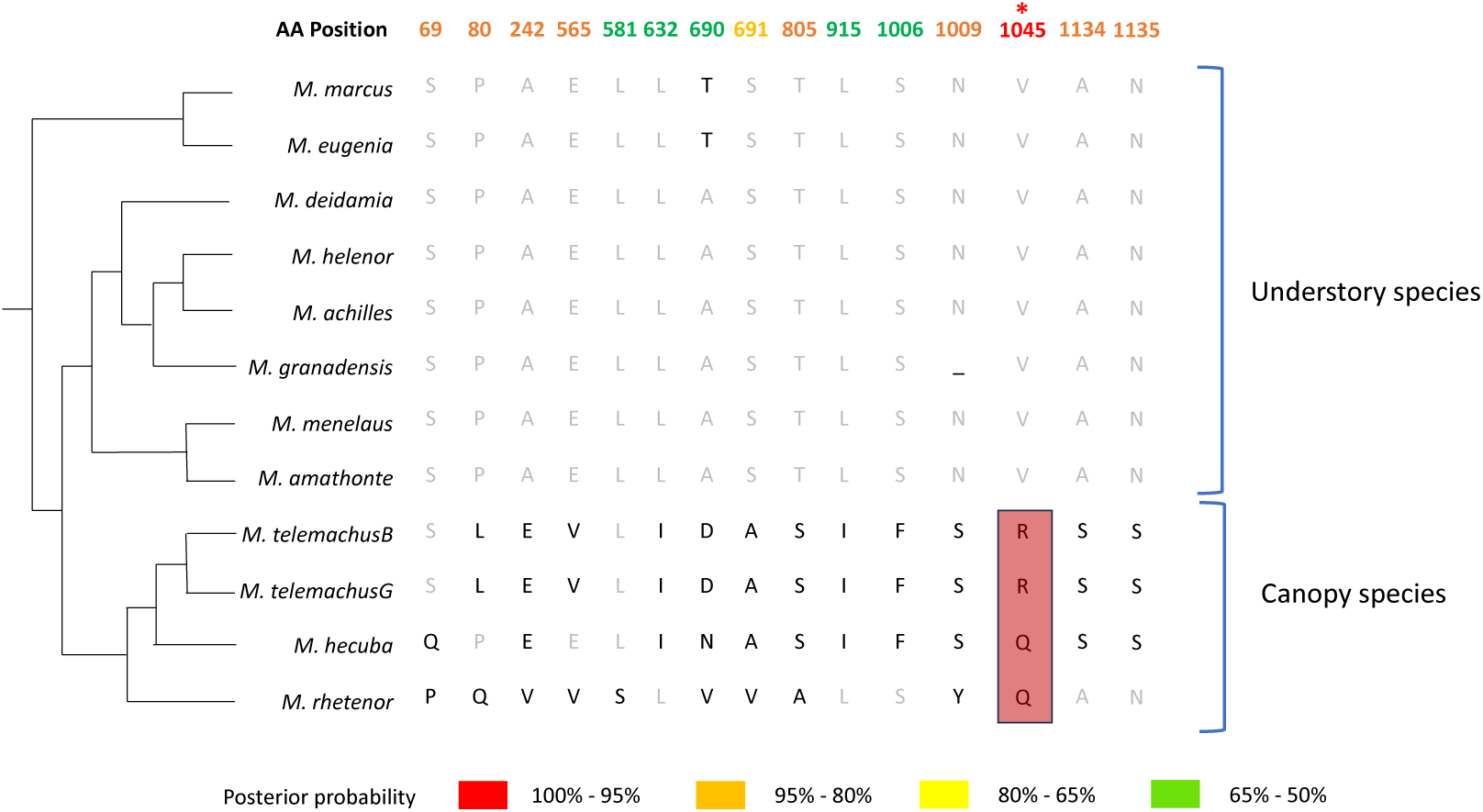
Partial amino acid alignment of the gene *Period* in *Morpho* species from the understory and the canopy, showing only the amino acids with a signature of positive selection using the branch site model in the PamL analysis. The position of these amino acids along the *Period* protein gene are indicated above in color. The values of the posterior probability associated with each site are indicated in colors, red indicating the highest probability (95 - 100%), while green showed the lowest probability (50 - 65%).

### Detection of selection at the genome scale

The *Pelican* analysis comparing gene evolution in the canopy *vs*. undestory species revealed a number of genes under positive selection (see the top-50 best candidate genes in table S3). The ontology of these genes did not reveal any functions that could be straightforwardly linked to adaptation to the different micro-habitats within the rainforest. We then investigated the location of these candidate genes on the different contigs, using the genome of *Morpho achilles* as a reference, to specifically test whether the Z chromosome was enriched in genes putatively under positive selection (table S3) In the list of the top 50 genes under positive selection in the canopy species, the top three contigs harboring the highest number of candidate genes were *ptg000021l* (6 candidate genes), *ptg000018l* (5 genes), and the Z-chromosome contig *ptg000022l* (5 genes). In the understory species, the top three contigs were the Z chromosome contig *ptg000022l* (8 genes), *pptg000020l* (5 genes) and *ptg000021l* (5 genes). In both micro-habitats, the Z-chromosome thus appears to contain a high number of genes under positive selection. This is unlikely to result from an elevated number of genes in this particular chromosome. In the genome of *M. achilles* (used here as a reference), the density of orthologs used for the *Pelican* analysis in the Z-chromosome contig was of 6.92 genes per Mb., while the contig *ptg000021l*, which also exhibited a large number of positively selected genes in both the canopy and understory species, showed a higher gene density (10.06 genes per Mb). The whole genome analysis of positive selection therefore reveals an elevated proportion of genes under divergent selection in the Z chromosome, when comparing species observed in the canopy vs. the understory, consistent with a significant effect of genes located in this sexual chromosomes in the ecological specialisation.

## Discussion

### Faster evolution of the Z-chromosome, likely driven by increased effects of drift and selection

By comparing the genomes of 11 *Morpho* species, our study revealed an increased evolution of the Z sexual chromosome relative to the autosomes in this genus: it exhibits a high propensity for genomic rearrangements and contains an elevated number of genes under positive selection.

Sex chromosomes have been observed to undergo accelerated rates of sequence evolution compared to autosomes [62], a phenomenon commonly known as Fast-X or Fast-Z evolution [68]. In birds for instance, the faster evolution of the Z-chromosome as compared to the autosomes has been evidenced in multiple species [35]. Faster evolution of sex chromosomes can result from genetic drift: for every mating pair, there are four copies of autosomal genes and three copies of X or Z-linked genes. Consequently, the effective population size (*Ne*) for X or Z sex chromosomes is three-quarters that of an autosome, intensifying the impact of genetic drift [36, 68]. Such an effect of drift was suggested to fuel the faster evolution of sex chromosomes observed in different groups with ZW system as some birds, snakes and the salmon louse *Lepeophtheirus salmonis* [45, 51, 69, 71] but see [18]. This enhanced drift could facilitate the fixation of different inversions in the Z chromosome of different *Morpho* species, as compared to the autosomes.

Faster rates of evolution in the X/Z chromosome is also enhanced by selection. The hemizygosity of X and Z chromosomes exposes rare advantageous recessive alleles to selection and leads to greater rates of fixation compared to rare autosomal alleles that are sheltered in heterozygotes [36]. Similarly, inversions located on the sex chromosomes are expected to disproportionately contribute to the evolution of local adaptation [16]. The greater proportion of inversions found on the Z-chromosome of *Morpho* butterflies could then be explained by the positive selective acting on the genes located within these rearrangements. Positive selection has been suggested to promote faster-X and Z evolution in mammals, *Drosophila*, but also in lepidoptera, like the silkmoth *Bombyx huttoni*, the moth *Manduca sexta* and the butterfly *Danaus plexippus* [48, 52, 62]. In the Crambidae moth *Ostrinia nubilalis*, a presumed 4 Mb inversion in the Z chromosome appears to increase the accumulation of ecologically adaptive alleles and genetic differentiation. Both the increased number of inversions and the higher number of gene under selection detected in the Z-chromosome in the *Morpho* genus could be fuelled by the divergent selective regime acting on the traits of *Morpho* species specialized in different micro-habitats.

### Ecological specialization generating divergent evolution on the Z-chromosome

The whole-genome analysis of signature of positive selection based on a model of amino acid sequence evolution, indeed showed that the Z chromosome is enriched in genes putatively under positive selection as compared to the autosomes, when comparing species specialised in the canopy and the understory habitats. The observed faster evolution of amino-acid sequences of genes located in the Z chromosome in our study aligns with findings in other lepidoptera species as the Carolina sphinx moth, the monarch butterfly or *Leptidea sinapis* assessed using dN/dS ratios [32, 52]. However, other studies in butterflies have failed to evidence a faster Z-chromosome evolution. In the species *Maniola jurtina* and *Pyronia tithonus*, transcriptome data showed a slightly slower evolutionary rate on the Z-chromosome compared to autosomes [61].

*Pelican*, that operates at the amino-acid level, has been shown to perform as well as other dN/dS commonly used methods to estimate positive selection at the genome level [21], however, some limitations in our analyses may impact our results. Our analyses were conducted on a subset of genes, specifically single-copy orthologs, identified through BUSCO and common across all genomes. BUSCO relies on OrthoDB, designed for multiple taxonomic lineages and comprising nearly universal groups of single-copy orthologs [75]. This subset, representing genes found in all Lepidoptera, may exclude genes with adaptive roles that are unique to the *Morpho* genus.

The specialization into different micro-habitats may have accelerated divergence in coding sequences in the sex-chromosome, as compared to the autosomes. Furthermore, reinforcement process might also favor divergent evolution of the Z-chromosome in sympatric species: heterospecific encounters in sympatry could enhance selection on genes favoring heterospecific avoidance, especially in females [44].

### Positive selection on a circadian gene located on the Z-chromosome

Since *Morpho* species living in sympatry display temporal segregation throughout the day [41], we specifically examined signatures of selection in various circadian genes previously documented in other Lepidoptera [49]. We found evidence of positive selection on specific sites within the *Period* gene, that is located on the Z chromosome in *Morpho*, as well as in all Lepidoptera species studied so far [64]. Among circadian genes, *Period* has been suggested to be involved in sympatric speciation across multiple taxa: variation in the amino-acid sequence of the *Period* protein, and variation in the regulatory regions of the *Period* gene have been shown to be involved in allochrony, *i.e.* differences in breeding time among conspecific individuals, that may facilitate speciation [66]. Experiments involving transgenic flies conducted by [65] indicated that *Period* played a role in speciation in both *D*. *melanogaster* and *D*. *pseudoobscura*, by exerting control over the precise timing of mating behavior [65]. In *Bactrocera cucurbitae*, *Period* was found to contribute to pre-mating isolation in two lines with distinct circadian periods, one short and one long [50].

In Lepidoptera, differences in diel activity are assumed to contribute to prezygotic reproductive isolation, as evidenced by the temporal segregation across 400 co-occurring species of Neotropical Hesperiidae [19]. Nevertheless, the role of circadian genes in regulating these differences or their putatively role in speciation has been scarcely studied, except in a few moth species. In *Spodoptera frugiperda*, the circadian gene *vrille* was found responsible for allochronic differentiation in the mating time of two different strains, by acting as a pre-mating barrier [29]. In the moth *Plutella xylostella*, the CRISPR/Cas9-mediated genome editing to knockout the *Period* gene changed circadian rhythms patterns in pupal eclosion, mating, egg-laying and egg hatching, suggesting that *Period* may be implicated in the timing of mating behaviour, as well as in other components of daily temporal niche in Lepidoptera [70]. The detection of a signal of positive selection on the Z-located gene *Period* in the genus *Morpho* could therefore be a consequence of the specialisation into different ecological niches, as a result of reinforcement. It may also suggest a role for *Period* in the speciation process, by acting as a pre-zygotic barrier.

### Z-chromosome inversions in sympatric species

Our findings reveal a higher prevalence of inversions in the Z chromosome as compared to autosomes, especially when comparing the genomes of sister-species coexisting in sympatry. Chromosomal inversions have been long acknowledged as a crucial mechanism that acts as a barrier to gene exchange, ultimately driving speciation [54]. In groups with a ZW system as birds, studies have shown an excess of inversions on the sex-chromosomes, and these inversions are more common between species with overlapping ranges than for species occurring in allopatry [33]. Because of their putative role as post-zygotic barriers, those inversions may have been selected after secondary contact, or have contributed to the initial speciation process. In the genus *Morpho*, we found that inversions on the Z chromosome were more common among closely-related species coexisting in sympatry, particularly between species with overlapping temporal niches, such as the canopy species *M* . *hecuba* and *M* . *telemachus*, or the understory sister-species *M* . *achilles* and *M* . *helenor* and *M* . *granadensis* and *M* . *helenor* [41][25]. Conversely, inversions were not observed in comparisons between closely related sympatric species with non-overlapping temporal niches, such as the basal sister-species *M* . *eugenia* (which flies very early in the morning around 6:00 AM) and *M* . *marcus* (which flies much later around 10:00 AM) [25], or between species found in allopatry, such as *M* . *amathonte* and *M* . *menelaus*. Our results are thus consistent with a putative role of inversions located on the Z-chromosome in the speciation and/or reinforcement process in the butterfly genus *Morpho*.

## Acknowledgments

We thank Pierre Lesturgie and Jean-Philippe Vernadet for help with scripts. We thank Louis Duchemin for all his help and assistance in using *Pelican*. All the analyses and simulations were performed on the the Plateforme de Calcul Intensif et Algorithmique PCIA (Muséum national d’histoire naturelle, Centre national de la recherche scientifique), the MeSU platform at Sorbonne-Université and the Genotoul bioinformatics platform Toulouse Occitanie (Bioinfo Genotoul, https://doi.org/10.15454/1.5572369328961167E12). This work was supported by the European Research Council (ERC) Consolidator Grant 101088089 “OUTOFTHEBLUE”

## Supporting Information

**Figure S1.**
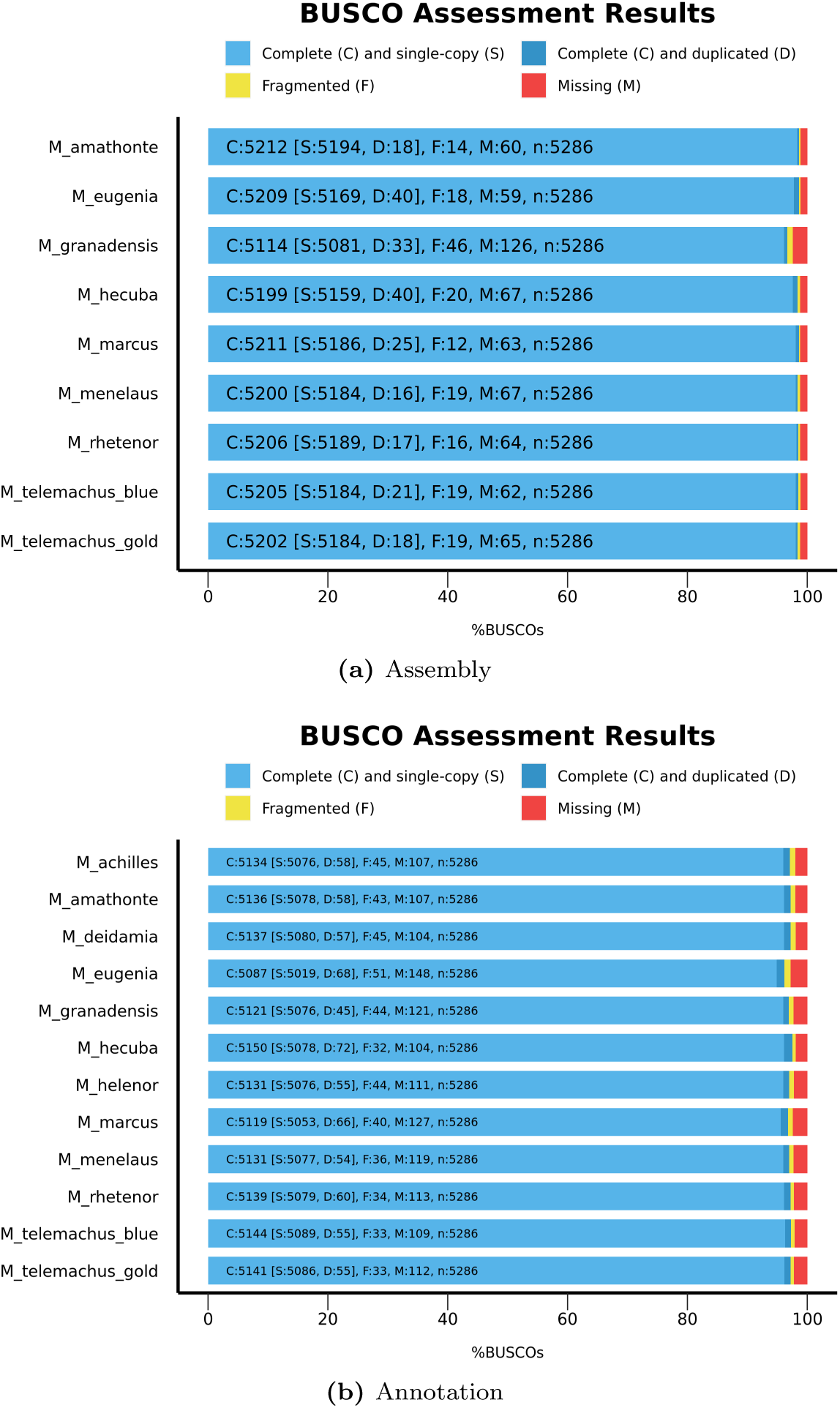
Quality assessment using BUSCO for above) assemblies (genome mode) and below) gene prediction (protein mode) for each *Morpho* species. Note that the BUSCO quality for the assemblies of *M* . *helenor*, *M* . *achilles* and *M* . *deidamia* have already been published and were then not included in the figure [3]

**Figure S2.**
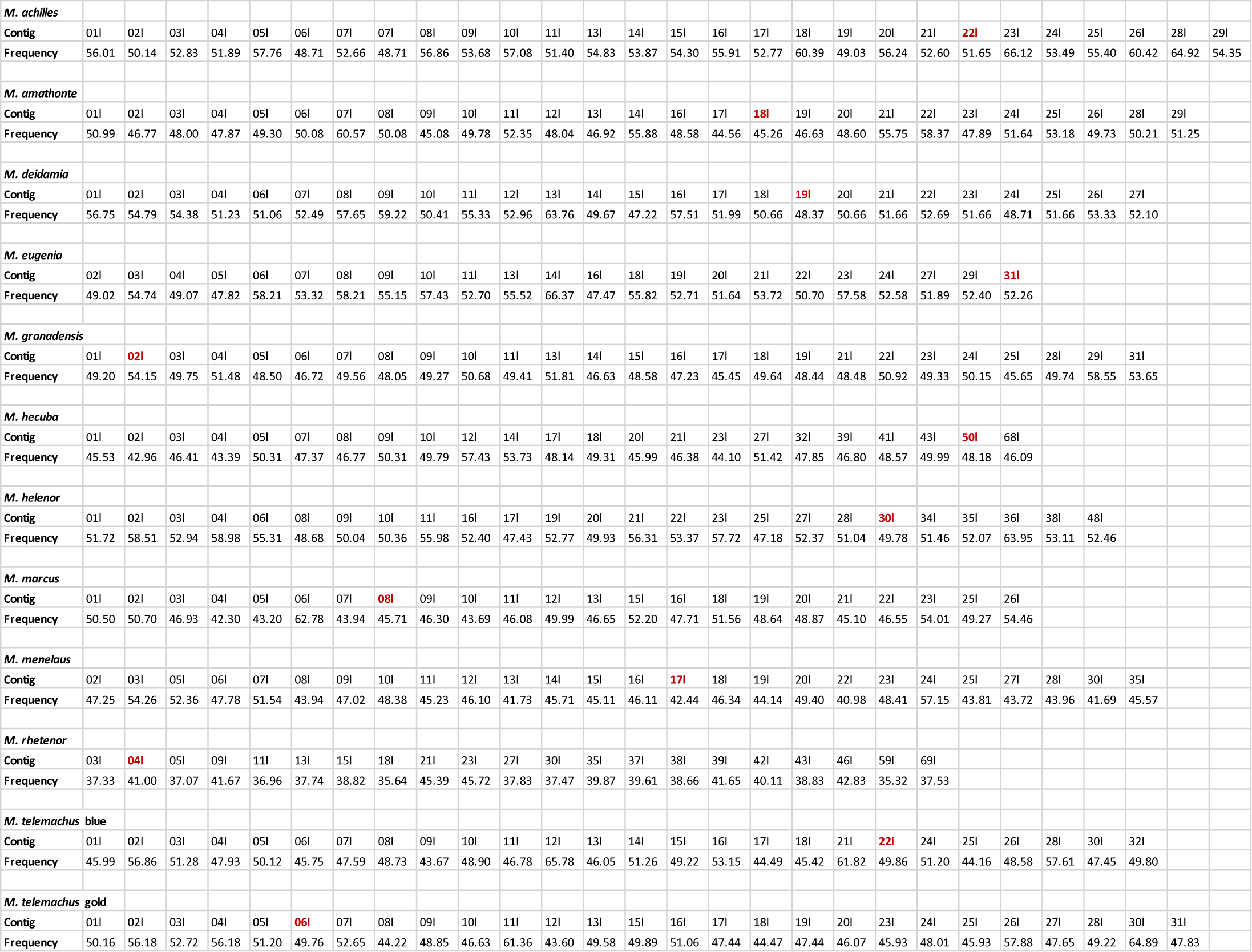
Percentage of repeated elements per contig (pt0000) across various *Morpho* genomes. For each species assembly, only contigs exceeding 10 megabytes were retained for the analyses. Contigs are uniquely identified by their names assigned by Hifiasm and are not consistent across different species. Additionally, the contig representing the Z chromosome is highlighted in red in all cases.

**Figure S3.**
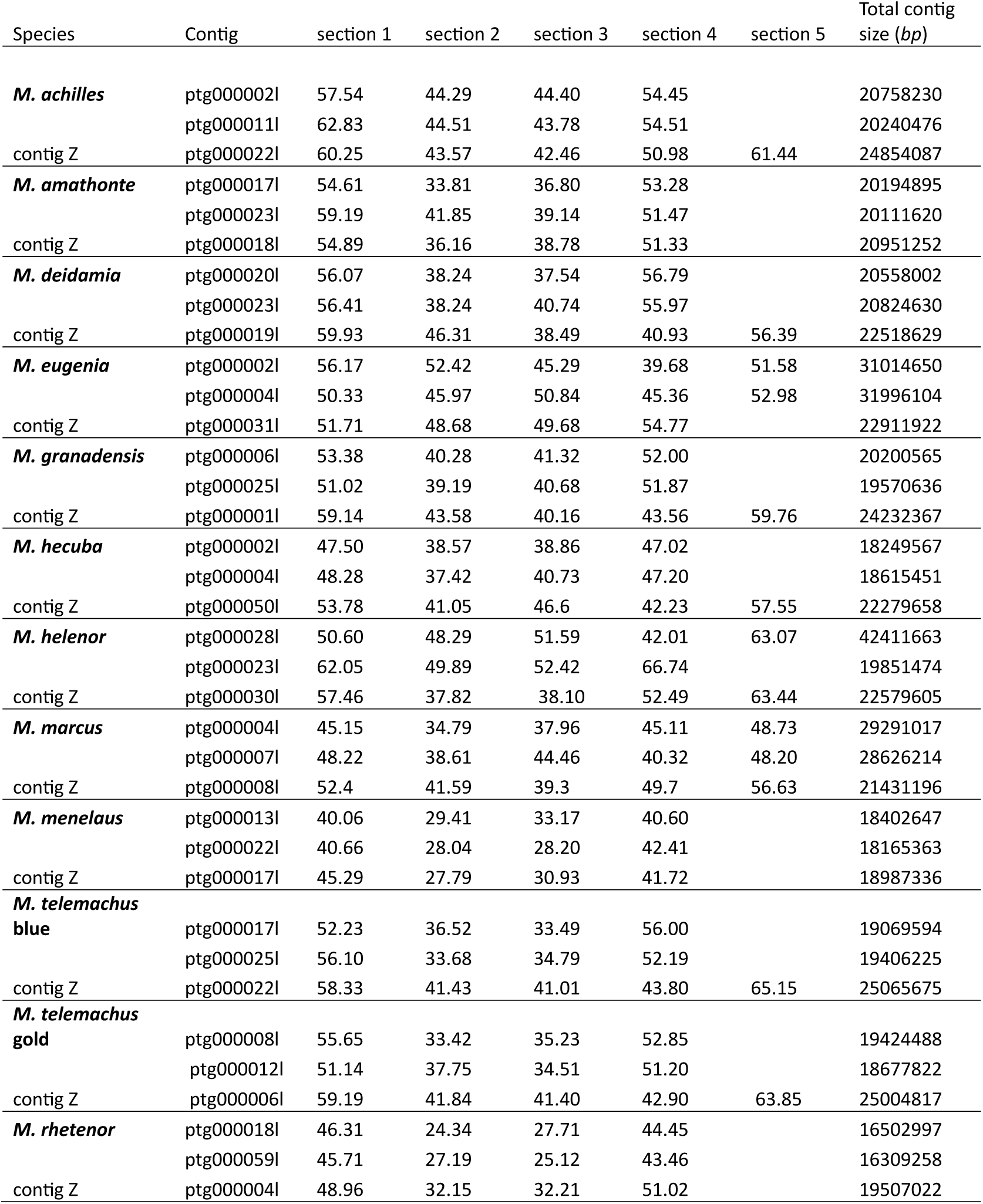
Frequency of repeated elements per section in the three biggest contigs (including the Z contig) of the assembly for every *Morpho* species. Each contig was partitioned into approximately 5 Mb sections, and RepeatModeler and RepeatMasker analyses were conducted for each section. The total size of the contig (in bp) is indicated in the last column.

**Figure S4.**
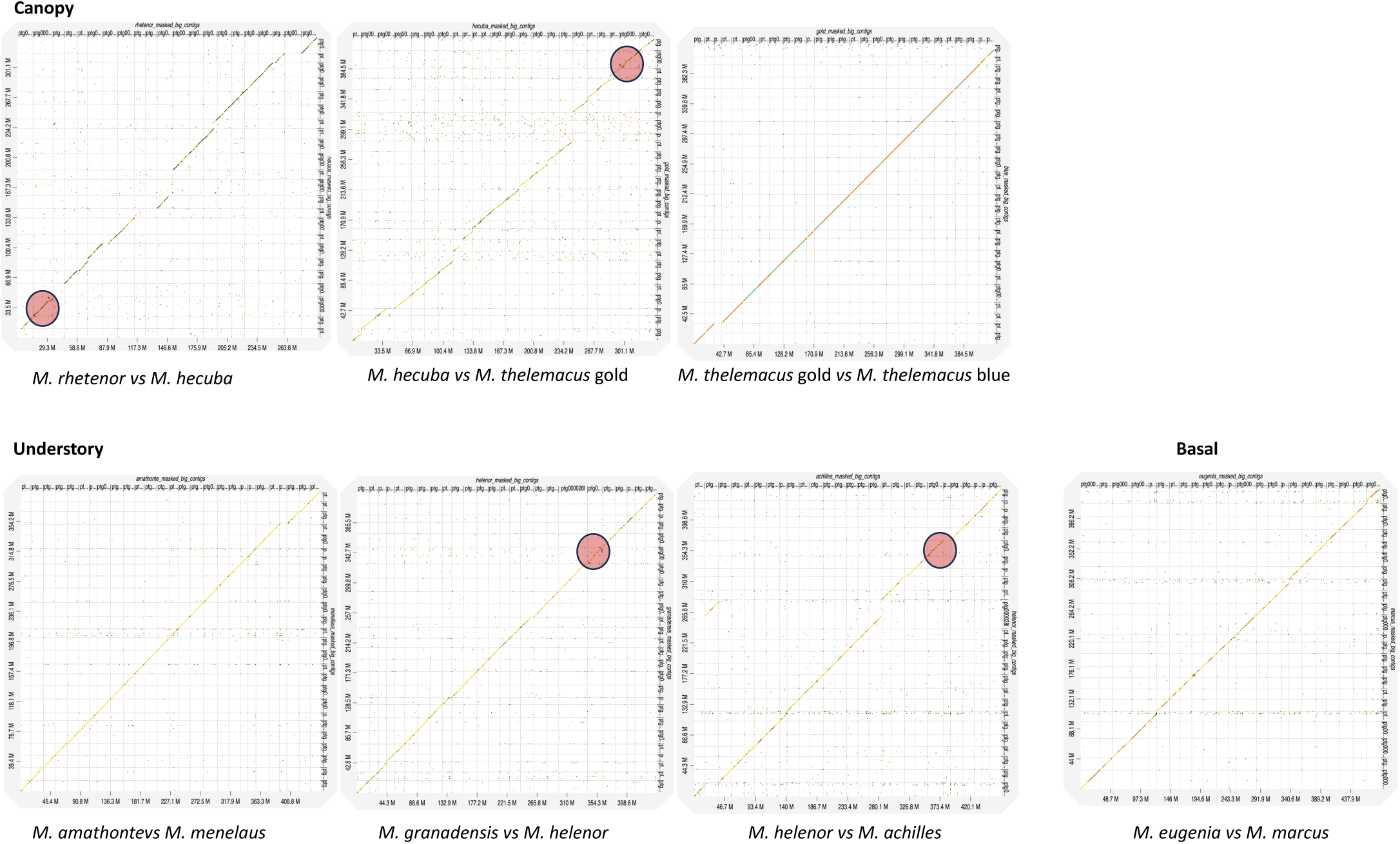
Paired whole-genome comparisons among closely related species of the genus *Morpho* belonging to three distinct clades: canopy, understory and basal. Genome assemblies were cleaned to retain only contigs of more than 10Mb to facilitate visualization. Detected genomic inversions were highlighted with red circles for ease of identification

**Figure S5.**
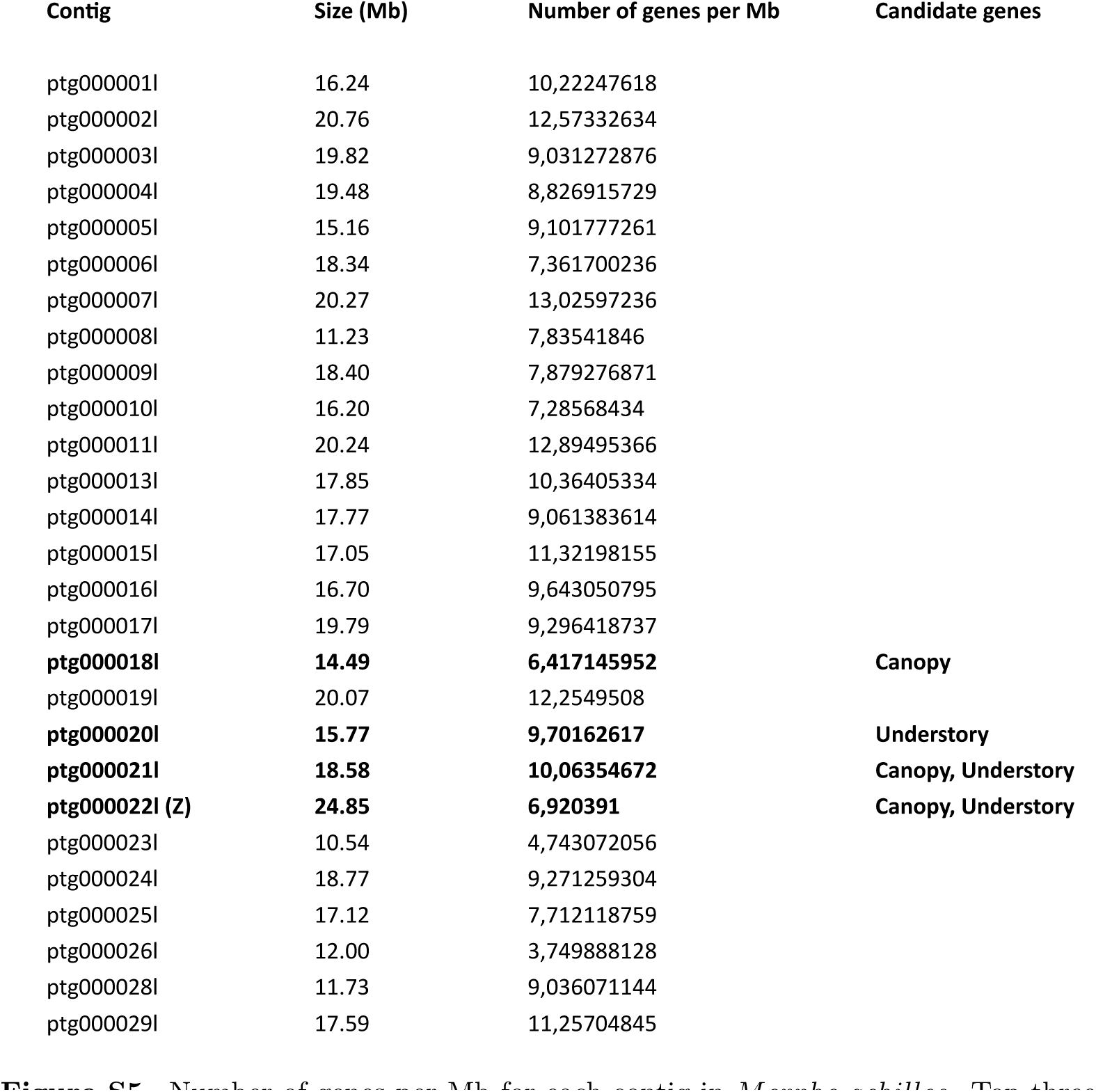
Number of genes per Mb for each contig in *Morpho achilles*. Top three contigs with the highest count of candidate genes among the top-50 candidates identified by *Pelican* analyses for the ’Canopy’ and ’Understory’ trees are marked in bold. Genes utilized in the analyses were sourced from BUSCO proteome results using busco2fasta.pl, representing single-copy orthologs present in all analyzed *Morpho* species (refer to the main text for details)

**FIG. S5.**
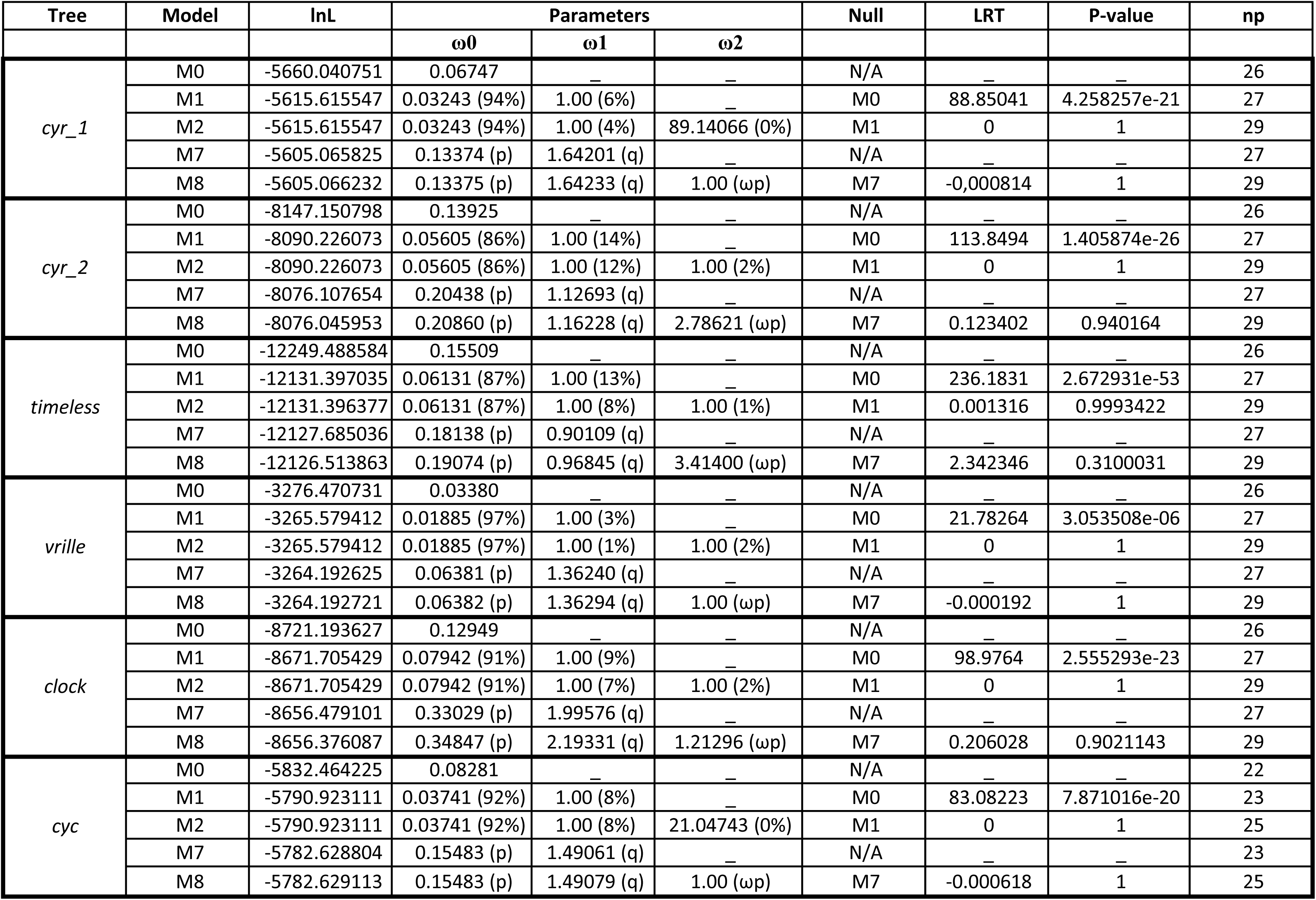
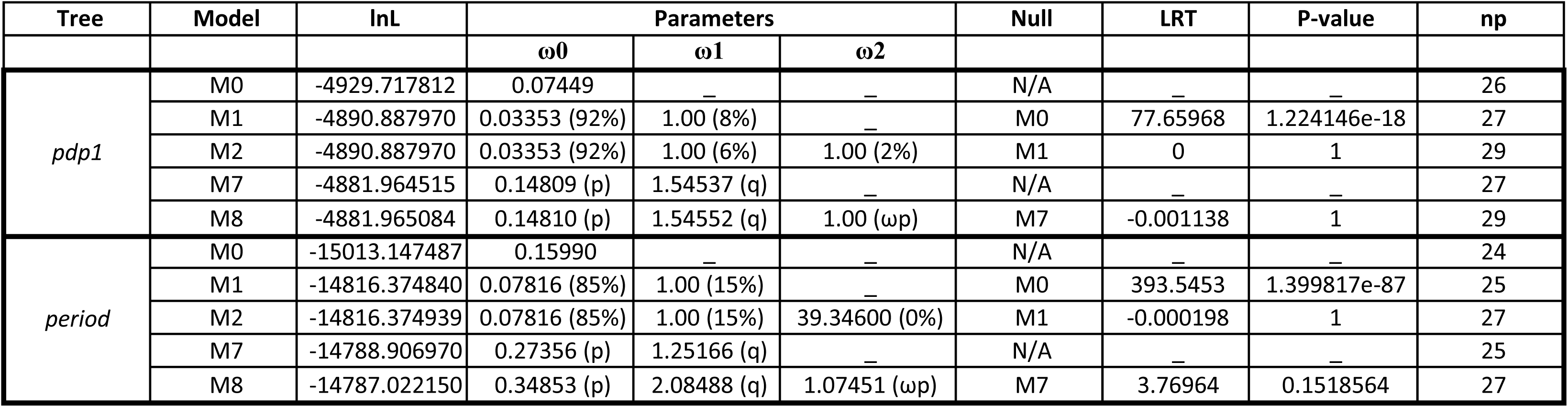
Results of PAML’s site-model analyses on eight circadian genes studied in the 11 *Morpho* species.

**FIG. S6.**
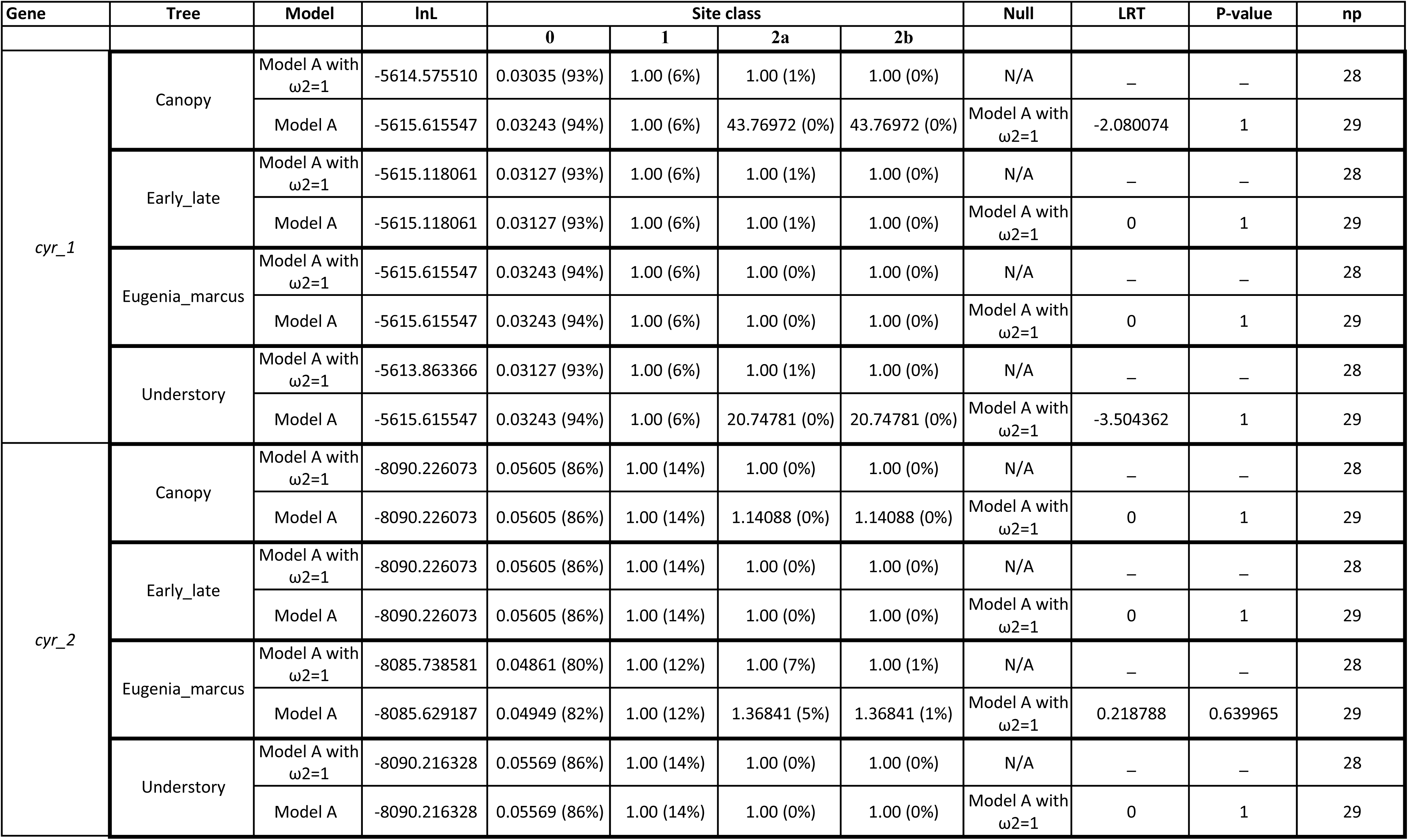
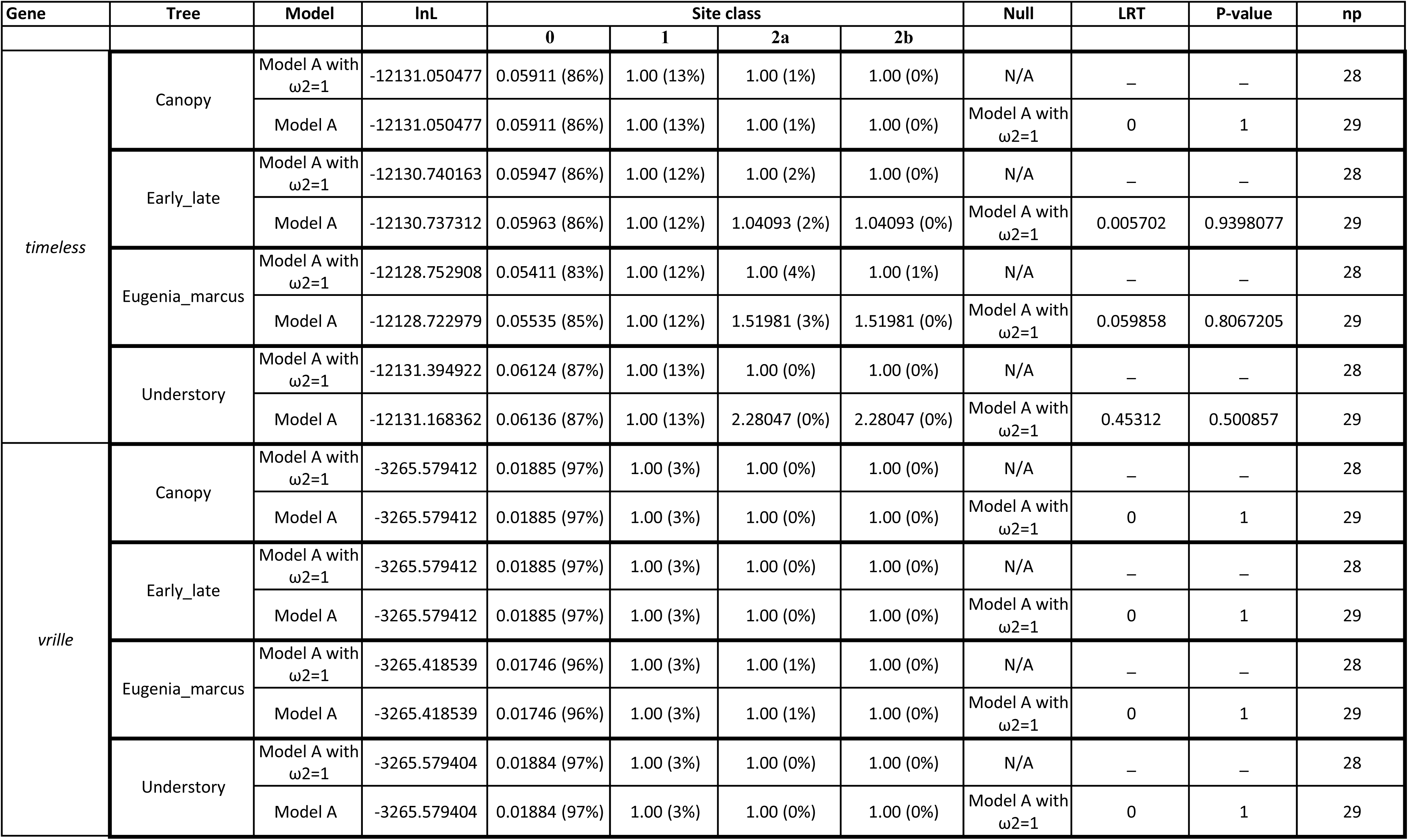
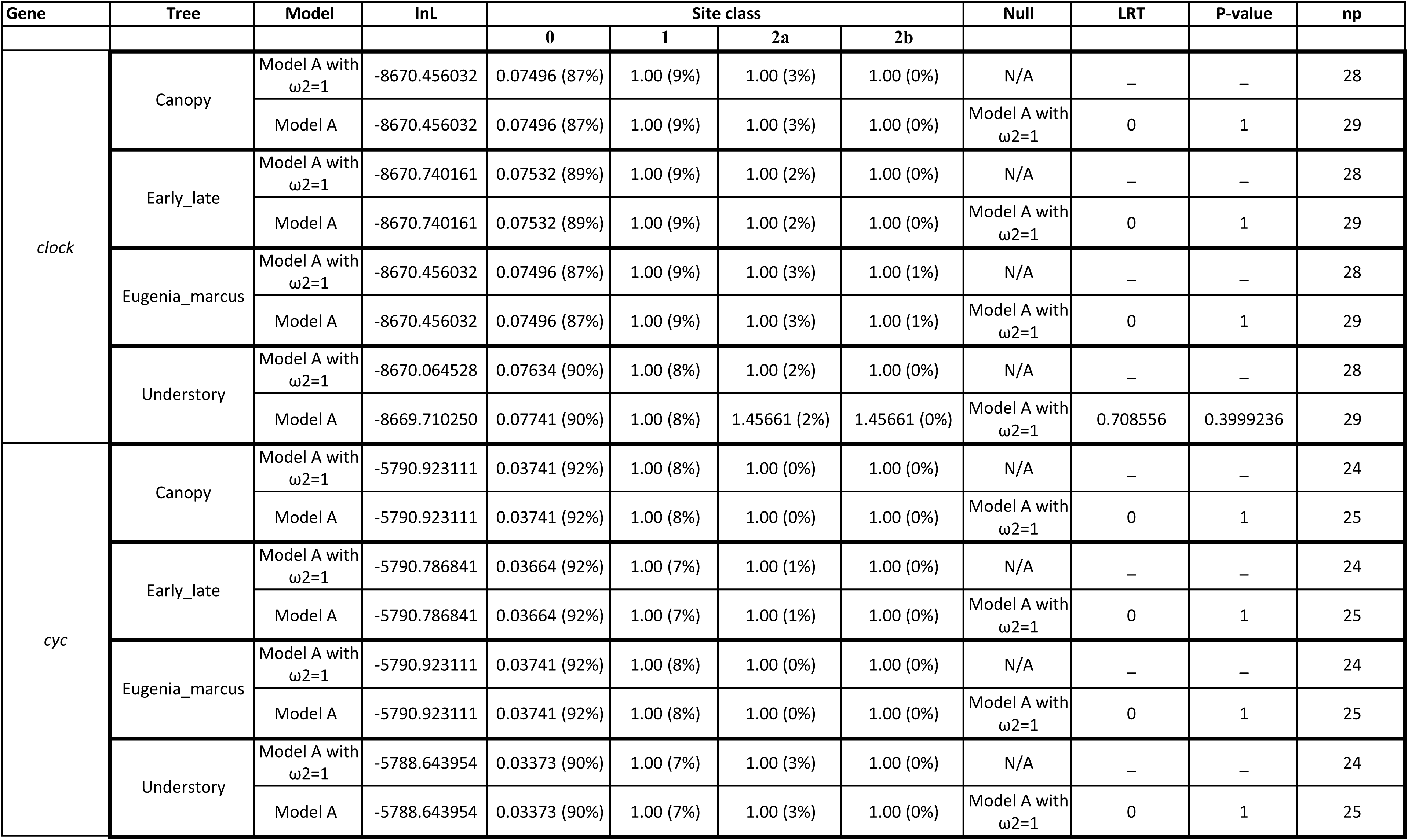
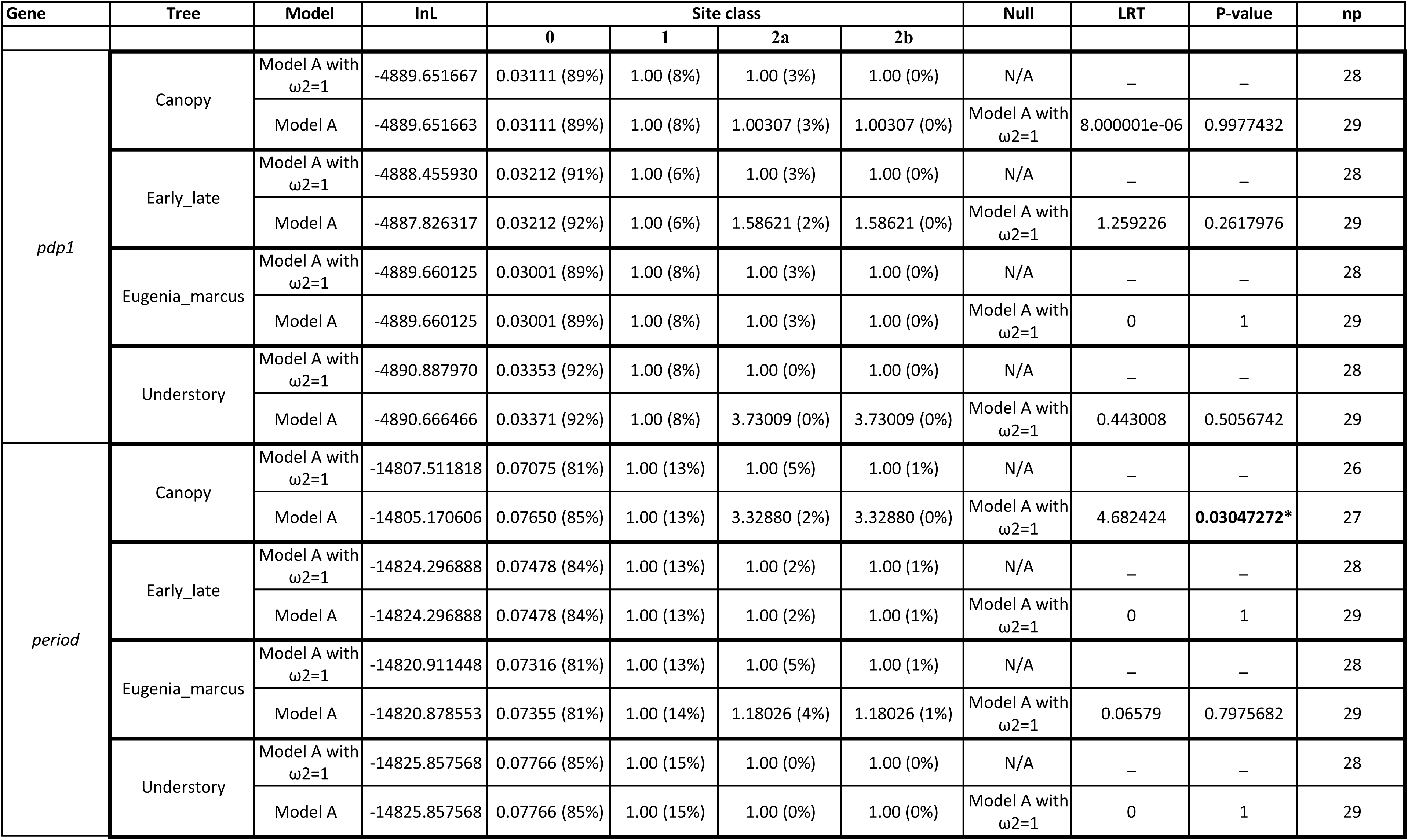
Results of PAML’s branch-site-model analyses on eight circadian genes in the 11 *Morpho* species. Bold values indicate significant p-values at the 0.05 significa

**Table S1.** Contig name and size of the Z chromosome in different *Morpho* species)

| Species | Contig | Size(Mb) |
| --- | --- | --- |
| *M. helenor* | ptg000030l | 22.58 |
| *M. achilles* | ptg000022l | 24.85 |
| *M. deidamia* | ptg000019l | 22.52 |
| *M. rhetenor* | ptg000004l | 19.51 |
| *M. hecuba* | ptg000050l | 22.28 |
| *M. amathonte* | ptg000018l | 20.95 |
| *M. menelaus* | ptg000017l | 18.99 |
| *M. granadensis* | ptg000001l | 24.23 |
| *M. eugenia* | ptg000031l | 22.91 |
| *M. marcus* | ptg000008l | 21.43 |
| *M. telemachus gold* | ptg000006l | 25.06 |
| *M. telemachus blue* | ptg000022l | 25.00 |

**Table S2.** Number of genes predicted for the different *Morpho* species using BRAKER2 and Funannotate (see text for details)

| Species | Number of genes |
| --- | --- |
| *M. helenor* | 29734 |
| *M. achilles* | 21017 |
| *M. deidamia* | 18620 |
| *M. rhetenor* | 18350 |
| *M. hecuba* | 20125 |
| *M. amathonte* | 20345 |
| *M. menelaus* | 19450 |
| *M. granadensis* | 18454 |
| *M. eugenia* | 20057 |
| *M. marcus* | 19239 |
| *M. telemachus gold* | 18300 |
| *M. telemachus blue* | 18159 |

**Table S3:**
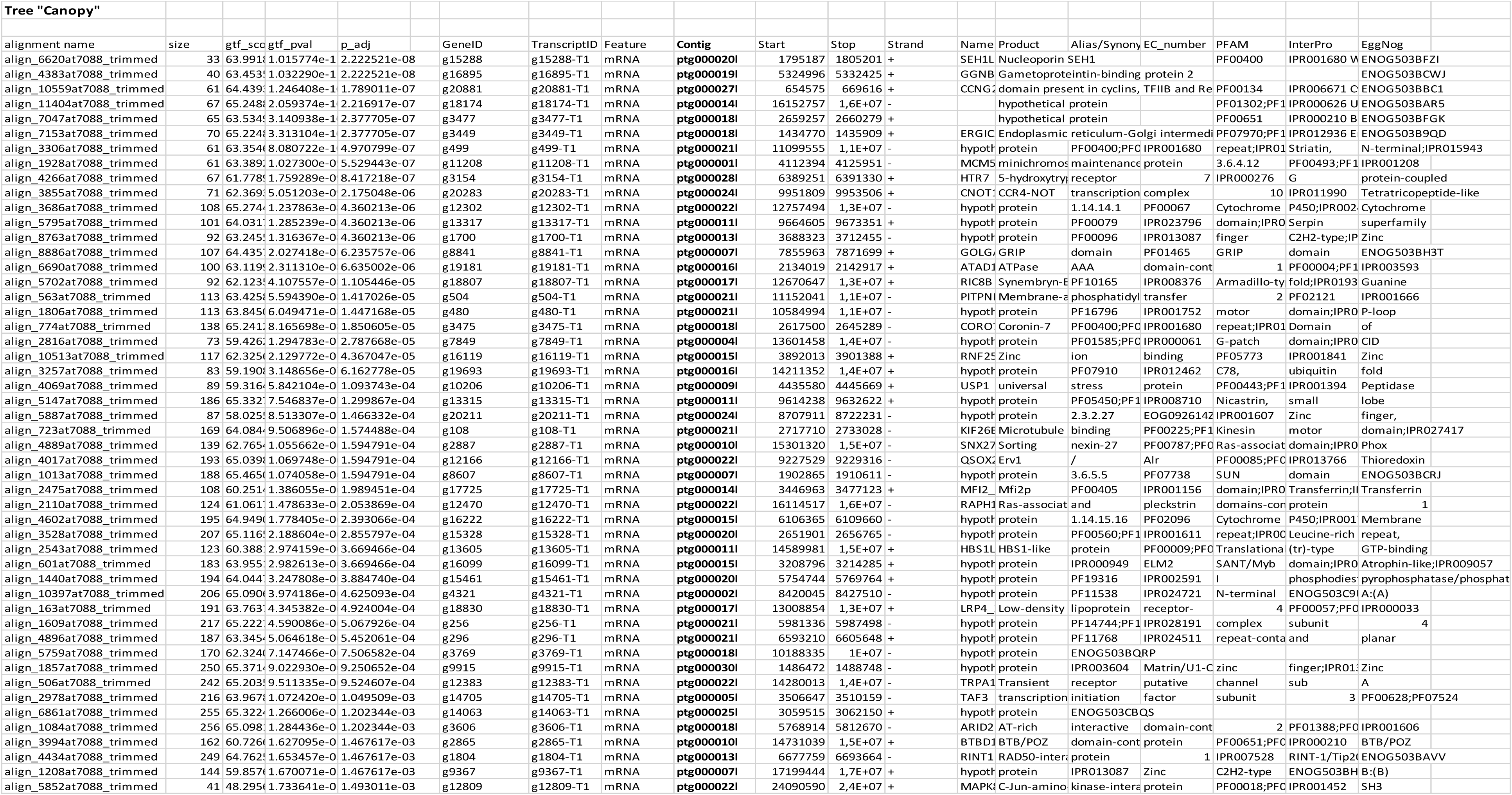
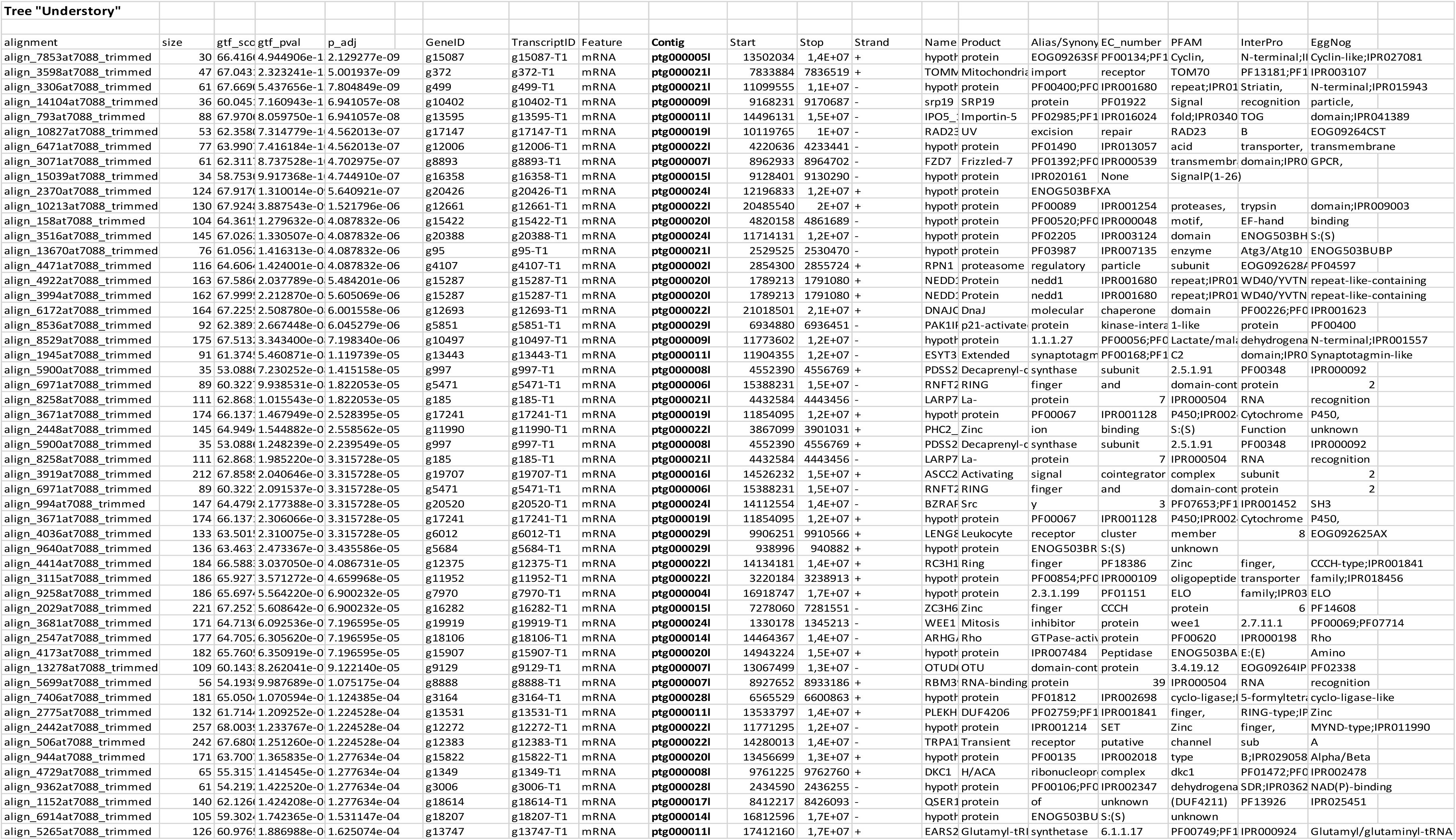
Top-50 best candidate genes from the analyses with *Pelican* for different trees: “Canopy” where the canopy species are treated in foreground (trait=1) and the understory species plus t labeled as background (trait=0) and “Understory” with the clade containing the understory species labelled as foreground and the canopy and basal species as background. The contig of location bold

